# Gene regulatory network analysis identifies sex-linked differences in colon cancer drug metabolism processes

**DOI:** 10.1101/277186

**Authors:** Camila M. Lopes-Ramos, Marieke L. Kuijjer, Shuji Ogino, Charles Fuchs, Dawn L. DeMeo, Kimberly Glass, John Quackenbush

## Abstract

Significant sex differences are observed in colon cancer, and understanding these differences is essential to advance disease prevention, diagnosis, and treatment. Males have a higher risk of developing colon cancer and a lower survival rate than women. However, the molecular features that drive these sex differences are poorly understood. We used both transcript-based and gene regulatory network methods to analyze RNA-Seq data from The Cancer Genome Atlas for 445 patients with colon cancer. We compared gene expression between tumors in men and women and found no significant sex differences except for sex-chromosome genes. We then inferred patient-specific gene regulatory networks, and found significant regulatory differences between males and females, with drug and xenobiotics metabolism via cytochrome P450 pathways more strongly targeted in females. This finding was validated in a dataset that included 1,193 patients from five independent studies. While targeting of the drug metabolism pathway did not change the overall survival for males treated with adjuvant chemotherapy, females with greater targeting had an increase in 10-year overall survival probability, with 89% (95% CI: 78%-100%) survival compared to 61% (95% CI: 45%-82%) for women with lower targeting, respectively (p=0.034). Our network analysis uncovered patterns of transcriptional regulation that differentiate male and female colon cancer. Most importantly, targeting of the drug metabolism pathway was predictive of survival in women who received adjuvant chemotherapy. This network-based approach can be used to investigate the molecular features that drive sex differences in other cancers and complex diseases.

## Introduction

Significant differences between the sexes are observed during the development and progression of diseases, influencing disease incidence and survival. For many cancer types, such as colon, skin, head and neck, esophagus, lung, and liver, males have a higher risk and higher mortality rates than females (1). Even though the higher risk in males might be attributed partially to occupational exposures and/or behavioral factors, such as diet, smoking and alcohol consumption, after adjusting for these risk factors males still have a higher cancer risk, although residual confounding cannot be excluded (1–3). In colon cancer, females not only have reduced risk relative to males, but also have a better prognosis (4–6). Furthermore, females have a higher survival benefit from 5-fluorouracil (5-FU)-based adjuvant chemotherapy as compared to males (7). Pharmacokinetics also vary between the sexes; females experience greater toxicity from certain chemotherapies, including 5-FU, consistent with the lower 5-FU clearance observed in females (8,9).

Sex differences in colon cancer have been largely attributed to sex hormones, yet the molecular mechanisms have not been established and clinical studies are contradictory (9). In general, studies point to the protective role of female hormones (estrogen) during colon cancer development and to the increased risk associated with male hormones (testosterone) (10–12). The circadian system might also explain the better prognosis in females, polymorphisms in the *CLOCK* sequence and the expression levels of miRNAs regulating the clock-genes were associated with longer overall survival of females with metastatic colorectal cancer compared to males (13). While previous studies focused on a few targeted genes, a systems-based analysis that integrates multi-omics data can provide insights into sex-specific regulatory processes associated with clinical outcome.

Regulatory networks characterize the complex cellular processes defined by a combination of signaling pathways and cell-type specific regulators. Each phenotype is defined by a characteristic network, while differences in network structures can shed light into the biological processes that distinguish phenotypes. Network-modeling approaches have been valuable in determining sex-specific regulatory features in healthy tissues and in disease (14,15).

While both the risk for, and outcome of colon cancer are different between men and women, clinical management is sex-independent. This may be because the molecular features that drive these sex differences are poorly understood. We used network-modeling approaches, PANDA (16) and LIONESS (16,17), to infer colon cancer patient-specific gene regulatory networks. We compared the male and female networks to identify genes that were targeted by transcription factors (TFs) in a sex-specific manner (Figure 1). We found that genes involved in drug and xenobiotics metabolism via cytochrome P450 were more strongly targeted by regulatory TFs in females; these results were validated in an independent dataset. Moreover, greater regulatory targeting of the drug metabolism pathway was predictive of longer survival in women who received adjuvant chemotherapy, but not in men.

**Figure 1.**
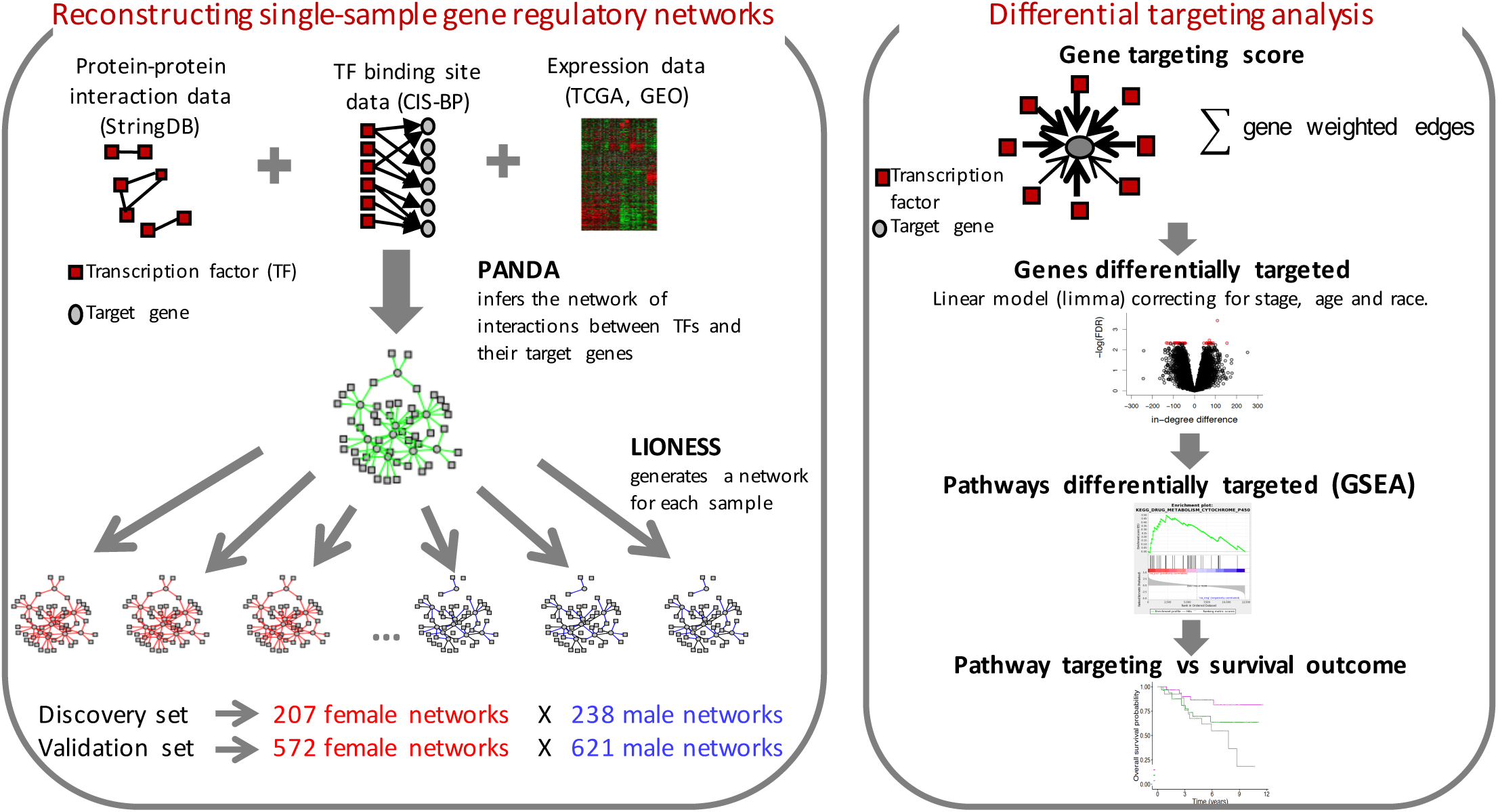
Study workflow. Overview of the approach used to reconstruct single-sample gene regulatory networks and to perform differential targeting analysis.

## Materials and Methods

### Discovery dataset

We downloaded level 3 RNASeqV2 and clinical data for colon cancer from The Cancer Genome Atlas (TCGA) on June 16, 2016 (https://tcga-data.nci.nih.gov). We kept only primary tumor samples, removed samples that were not annotated for sex, and used principal component analysis (using the plotOrd function in metagenomeSeq 1.12.1 (18)) on genes located on the Y chromosome to identify and remove 9 potential sex-misannotated samples. After performing these quality control steps, the discovery dataset included 445 primary colon tumor samples before treatment, from 238 males and 207 females.

We filtered lowly expressed genes by removing genes with less than one count per million (CPM) in at least 104 samples, using the cpm function from R package edgeR 3.18.1 (19), which corresponded to 5,571 out of 20,249 genes. We chose 104 samples because that represents half of the samples in the smaller subgroup. To retain the same set of genes in the discovery and validation datasets, and the same set of genes for the differential expression and differential targeting analysis, we kept only the genes overlapping the filtered genes in the discovery dataset, the validation dataset, and the genes in the TF/target gene regulatory prior used for creating the gene regulatory networks (see sections: “Validation dataset” and “Single-sample gene regulatory networks and differential targeting analysis”). This corresponded to 12,817 genes, which included genes on the sex chromosomes.

The expression data generated by TCGA were normalized using smooth quantile normalization (20), and batch was corrected for sequencing platforms (IlluminaGA and IlluminaHiSeq) and shipment date, as implemented in the R package qsmooth available on Github (https://github.com/stephaniehicks/qsmooth).

### Validation dataset

We searched the Gene Expression Omnibus (GEO) repository for colon cancer studies that included patient survival data, and were obtained from the same microarray platform (Affymetrix Human Genome U133 Plus 2.0 Array). The validation dataset set contained five independent studies: GSE14333, GSE17538, GSE33113, GSE37892, and GSE39582. Raw expression data and clinical data were downloaded from GEO using the R package GEOquery 2.36.0 on July 10, 2015. Raw expression data were pre-processed by frozen robust multiarray analysis (fRMA) using the R package frma 1.22.0 (21). Genes with multiple probe sets were represented by the probe set with the highest mean across all datasets. Then, we only kept the genes that overlapped with the discovery dataset, and the genes in the TF/target gene regulatory prior. We removed samples obtained from rectal tumor, samples that were not annotated for sex, and potential sex-misannotated samples (n=84), as previously described. We performed batch correction by dataset series using the ComBat function implemented in the R package sva 3.18.0 (22). The final validation dataset was comprised of 1,193 primary colon tumor samples before treatment, from 621 males and 572 females, and 12,817 genes.

### Differential expression analysis

We used voom available in the R package limma 3.26.9 (23) to compare gene expression between colon tumor samples from males and females after adjusting for the covariates age, race, and disease stage by the Union for International Cancer Control (UICC). We performed multiple testing corrections using Benjamini-Hochberg (24).

### Single-sample gene regulatory networks and differential targeting analysis

We used the PANDA (16) and LIONESS (17) algorithms to reconstruct gene regulatory networks for each sample in both the discovery and the validation datasets. The networks were inferred from three types of data: TF/target gene regulatory prior (obtained by mapping TF motifs from the Catalog of Inferred Sequence Binding Preferences (CIS-BP) (25) to the promoter of their putative target genes), protein-protein interaction data (using the interaction scores from StringDb v10 (26) between all TFs in the regulatory prior), and gene expression (obtained from the discovery or validation datasets). The TF/target gene regulatory prior, and the protein-protein interaction data were generated as described by Sonawane et al. (27), and then we kept only the genes that matched the genes expressed in the discovery and validation datasets. Our TF/target gene regulatory prior consisted of 661 TFs targeting 12,817 genes, and the protein-protein interaction data consisted of interactions between the 661 TFs.

For each sample’s gene regulatory network, we calculated each gene targeting score, equivalent to its gene’s in-degree (defined as the sum of all incoming edge weights from all TFs in the network). Next, we compared the gene targeting score between males and females using a linear regression model and correcting for the covariates age, race, and disease stage, as available in the R package limma 3.26.9 (28).

The subnetwork was illustrated using Cytoscape default yFiles Organic layout (version 3.4.0) (29) where each edge connects a TF to a target gene, and the color represents the average edge weight difference between the male and female networks.

### Pathway enrichment analysis

To perform pathway enrichment analysis, we used pre-ranked Gene Set Enrichment Analysis (GSEA) (Java command line version 2-2.0.13) (30), and the gene sets obtained from the Kyoto Encyclopedia of Genes and Genomes (KEGG) pathway database (31) that were downloaded from the Molecular Signatures Database (MSigDB) (http://www.broadinstitute.org/gsea/msigdb/collections.jsp) (“c2.cp.kegg.v5.0.symbols.gmt”). We only considered gene sets of size greater than 15 and less than 500 after filtering out those genes not in the expression dataset, which restricted our analysis to 176 gene sets. To perform the analysis, all genes were ranked by the *t*-statistic produced by the voom differential expression analysis or by the limma differential targeting analysis after adjusting for covariates.

### Survival analysis

We performed Kaplan–Meier survival analysis as implemented in the R package survival 2.41-3, and the p-values were computed using the log-rank test. The survival curves were plotted using the function ggsurv on the GGally package 1.3.2.

## Results

### Data features

We investigated the regulatory processes that drive sex differences in colon cancer by performing network analysis in a discovery dataset followed by a validation analysis in an independent dataset. For the discovery dataset, we analyzed RNA-Seq data of primary colon tumor samples from TCGA. We removed potential sex mis-annotated samples after assessing the expression of Y chromosome genes, and retained 445 samples, obtained before treatment, for the primary analysis.

For the validation dataset, we downloaded raw microarray data from five independent studies in GEO (32), which were profiled on the same microarray platform (Affymetrix Human Genome U133 Plus 2.0 Array). We corrected the data for study batch, and removed potential sex mis-annotated samples. The final validation dataset included 1,193 primary colon tumor samples obtained before treatment. Patient clinical features are summarized in Table 1 and are extensively presented in Table S1. The number of males and females were well matched, and for all downstream analysis we controlled for potential differences between the sexes for age, race, and colon cancer stage.

**Table 1.**
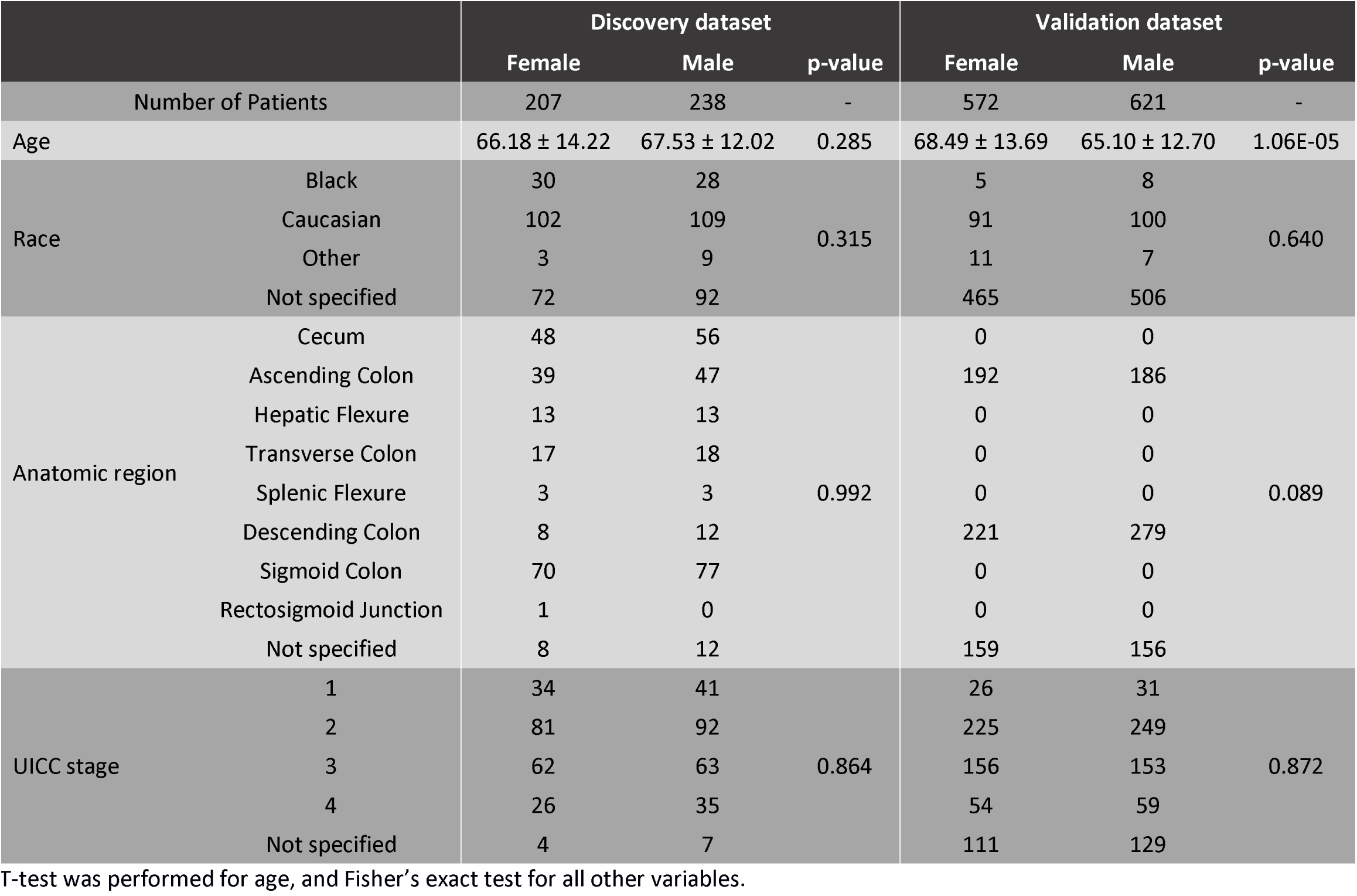
Clinical features for the discovery (TCGA) and validation (GEO) datasets included in the study.

### Autosomal genes are not differentially expressed between males and females in colon cancer

As a baseline for our network analysis, we first performed a differential expression comparison between males and females using voom (23). We found twelve genes significantly different between males and females with an absolute fold change greater than two and a false discovery rate (FDR) less than 0.1 (Figure 2A). All the differentially expressed genes were located on the X and Y chromosomes, and no genes were found to be differentially expressed when we excluded sex chromosomes from the analysis.

**Figure 2.**
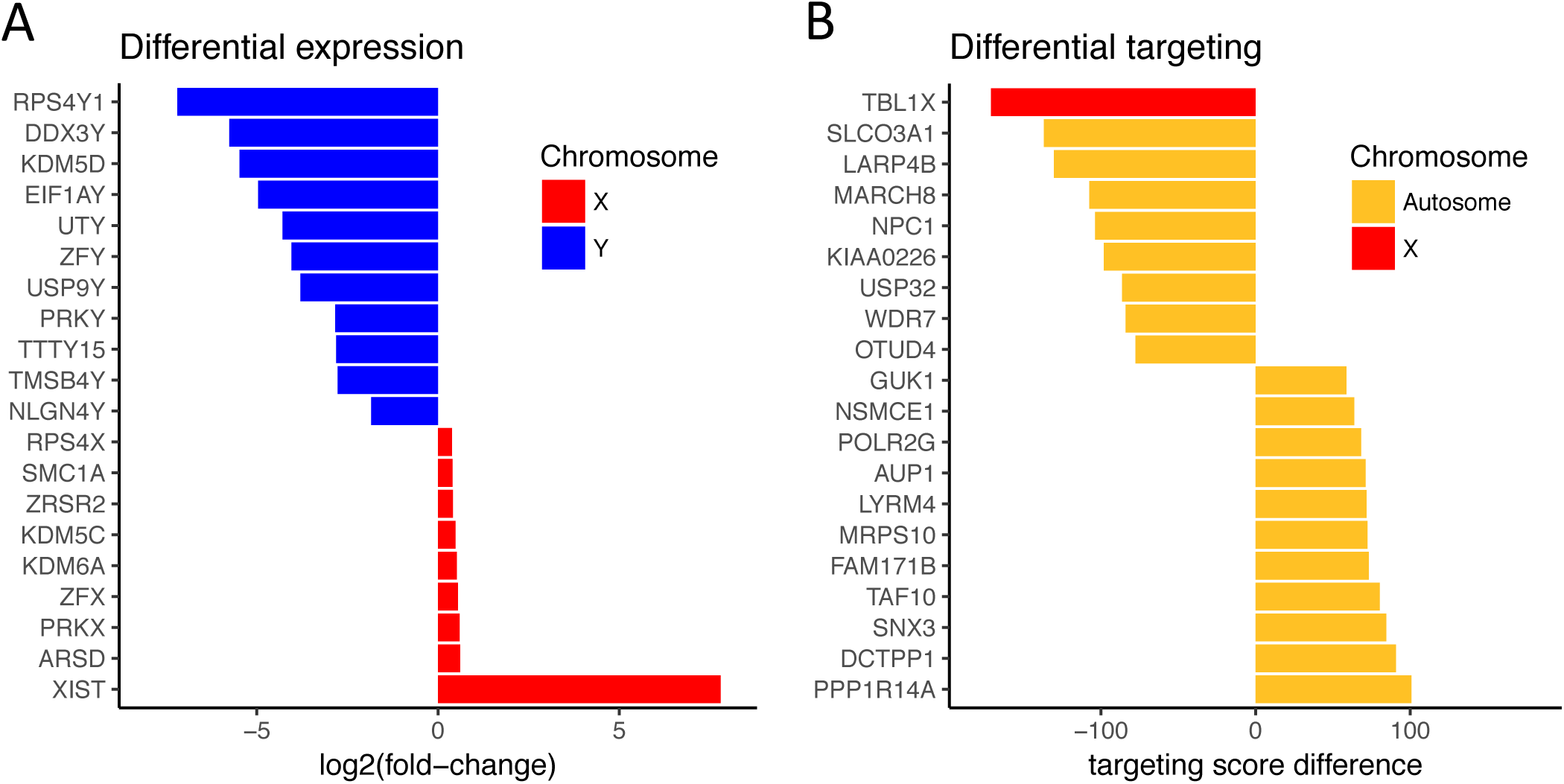
Differential expression and differential targeting between male and female colon cancer. A) Expression log_2_(fold-change) of the top 20 differentially expressed genes between males and females in the discovery dataset. B) Gene targeting score difference of the top 20 genes differentially targeted between males and females in the discovery dataset. Positive values indicate higher levels in females, and negative values indicate higher levels in males.

We also tested whether there were changes in gene expression levels across the genome that were below our detection threshold but which might be relevant for sex-differences in colon cancer. Therefore, to understand the biological functions associated with changes in gene expression, we ranked all genes based on their statistical significance (t-statistic) and performed pre-ranked GSEA using KEGG pathways (30,31). We did not find any significant differential enrichment of KEGG pathways between males and females. Thus, based only on gene expression differences between the sexes, it was not possible to identify biological pathways that distinguish males and females with colon cancer.

### Differential targeting of biological pathways in males and females

In previous studies, we have found gene regulatory network analysis provides greater insight into altered biological processes than do simple tests of differential gene expression (15,27,33). Thus, we performed a gene regulatory network analysis using PANDA (16) and LIONESS (17) (Figure 1), inferring gene regulatory networks for each individual in our discovery population and independently for the validation dataset.

PANDA is a message-passing algorithm that starts with a TF/target gene prior regulatory network based on a motif scan that maps TFs binding sites to the promoter of their putative target genes, then integrates the prior regulatory network with protein-protein interaction data, and gene expression data. In the final gene regulatory network, each edge connects a TF to a target gene, and the associated edge weight reflects the strength of the inferred regulatory relationship. LIONESS uses an iterative process that leaves out each individual in a population, estimates the network with and without that individual, and then interpolates edge weights to derive an estimate for the network active in that single individual. This method enables identifying individual variability in the regulatory network, and allows us to associate network properties with clinical information. We initiated PANDA with TF binding sites from CISBP (25), protein-protein interaction data from StringDb (26), and expression data from the discovery or validation sets to estimate the aggregate gene regulatory network based on all samples. Then we used LIONESS to estimate each sample-specific gene regulatory network.

As a control for the differential targeting analysis, we started by comparing colon cancer with healthy colon tissue. For the healthy colon networks, we used the single-sample gene regulatory networks reconstructed by Chen et. al. (34), which were modeled on data from the Genotype-Tissue Expression (GTEx) project (35). For the colon cancer networks, we reconstructed gene regulatory networks for each sample as shown in Figure 1. Next, we performed differential targeting analysis. For each gene, we calculated a gene targeting score equal to the sum of all incoming edge weights a gene receives from all TFs in the network (its “in-degree”). We then used a linear regression model to test whether there is a significant difference in the gene targeting score between healthy and cancer tissues using limma (24), and adjusting for age, and race. All genes were ranked by their statistical significance, and we used pre-ranked GSEA to identify the pathways enriched for the differentially targeted genes. As expected when comparing healthy and cancer tissues, we found that pathways enriched for genes more strongly targeted in cancer were associated with apoptosis and immune signaling, while pathways enriched for genes more strongly targeted in healthy tissues were associated with focal adhesion and extracellular matrix interaction (Figure S1). Similar pathways were found to be differentially targeted between healthy and cancer tissues independent of the sex analyzed.

To explore sex differences in colon cancer, we performed a similar differential targeting analysis to compare the male colon cancer networks with the female colon cancer networks and identified the genes differentially targeted between the sexes. We found 21 genes differentially targeted between males and females (FDR<0.1); only one of the genes (*TBL1X*) is located on sex chromosomes (Figure 2B). This stands in contrast to the differential expression analysis which found genes exclusively on the sex chromosomes. While the differential expression analysis gives the expression difference for each individual gene, the differential targeting analysis allows us to infer how much a gene is targeted by a set of TFs, and may better elucidate the biological differences in colon cancer between males and females.

We ranked all genes based on their differential targeting statistical significance (t-statistic), and performed pre-ranked GSEA. We found that genes more strongly targeted in males were enriched for NOTCH signaling pathway (FDR=0.002), mTOR signaling pathway (FDR=0.01), and WNT signaling pathway (FDR=0.02). These pathways have key roles in colon cancer biology (36,37), and their higher targeting in males may be related with the higher risk and lower survival rates observed in colon cancer in males. In females, we found significantly higher targeting of pathways associated with metabolism, including steroid hormone biosynthesis (FDR=0.02), metabolism of xenobiotics by cytochrome P450 (FDR=0.02), and drug metabolism-cytochrome P450 (FDR=0.04) (Table 2). This indicates that regulatory differences between the sexes in the drug metabolism pathway may impact the response to chemotherapy treatment and survival outcome in a sex-specific manner. We therefore further investigated this pathway.

**Table 2.** Pathways differentially targeted between male and female colon cancer regulatory networks.

| Enrichment in males |  |  | Enrichment in females |  |  |
| --- | --- | --- | --- | --- | --- |
| Pathway | NES | FDR | Pathway | NES | FDR |
| Acute myeloid leukemia | -2.207 | 0.000 | Oxidative phosphorylation | 2.637 | 0.000 |
| Endometrial cancer | -2.230 | 0.000 | Parkinsons disease | 2.485 | 0.000 |
| Chronic myeloid leukemia | -2.157 | 0.001 | Ribosome | 2.287 | 0.000 |
| Notch signaling pathway | -2.114 | 0.002 | Alzheimers disease | 1.957 | 0.003 |
| Phosphatidylinositol signaling system | -2.062 | 0.002 | Proteasome | 1.968 | 0.004 |
| ErbB signaling pathway | -2.006 | 0.002 | Peroxisome | 1.914 | 0.006 |
| Pancreatic cancer | -2.012 | 0.003 | Huntingtons disease | 1.872 | 0.008 |
| Focal adhesion | -2.018 | 0.003 | Terpenoid backbone biosynthesis | 1.807 | 0.015 |
| Colorectal cancer | -2.026 | 0.003 | Metabolism of xenobiotics by cytochrome p450 | 1.791 | 0.015 |
| Prostate cancer | -1.962 | 0.003 | Steroid hormone biosynthesis | 1.761 | 0.018 |
| Jak stat signaling pathway | -1.923 | 0.004 | Protein export | 1.734 | 0.021 |
| Adherens junction | -1.914 | 0.004 | Histidine metabolism | 1.736 | 0.022 |
| Arrhythmogenic right ventricular cardiomyopathy arvc | -1.917 | 0.004 | Amyotrophic lateral sclerosis als | 1.677 | 0.034 |
| Fc gamma r mediated phagocytosis | -1.886 | 0.005 | Drug metabolism cytochrome p450 | 1.648 | 0.041 |
| Chemokine signaling pathway | -1.849 | 0.007 | Tyrosine metabolism | 1.590 | 0.064 |

NES: Normalized enrichment score, FDR: False discovery rate

### Genes involved in drug metabolism are more strongly targeted by transcription factors in females

The network analysis identified significant sex-differences in the regulatory pattern of the drug metabolism pathway. Since the regulatory differences are present in non-treated tumor tissues, these differences can set the basis on how each sex will respond to chemotherapy treatment, and impact sex-specific management and outcome.

We compared the edge weights connecting TFs with genes using limma (correcting for age, race, and stage). Figure 3A illustrates the edges around genes in the drug metabolism pathway that are most significantly different between males and females. Of the 2,830 edges around genes in the drug metabolism pathway, 220 edges have a FDR<0.05 and are represented in Figure 3A. Overall, we observe most significant edges are those with strong regulatory targeting of genes in the female networks, contributing to the higher gene targeting score we had found. Many of the genes more strongly targeted in females, such as *GSTO1, GSTA4, GSTT2, MGST2, MGST3*, belong to the family of glutathione S-transferases, which are important in the detoxification leading to the elimination of toxic compounds (38), whereas in the male networks only *GSTM2* is more strongly targeted. We also find that some genes, such as *CYP2D6* and *GSTM4*, have similar overall gene targeting score, while being targeted by different sets of TFs in male and female networks. For example, in the female networks, *CYP2D6* is more strongly targeted by TFs responsive to estrogens, such as estrogen related receptor alpha (*ESRRA*), estrogen related receptor beta (*ESRRB*), and estrogen related receptor gamma (*ESRRG*). In the males networks, *CYP2D6* is more strongly targeted by SRY-box 12 (*SOX12*), which belongs to a family of TFs characterized by the presence of a DNA-binding high mobility group domain, homologous to that of sex-determining region Y (*SRY*) (39).

**Figure 3.**
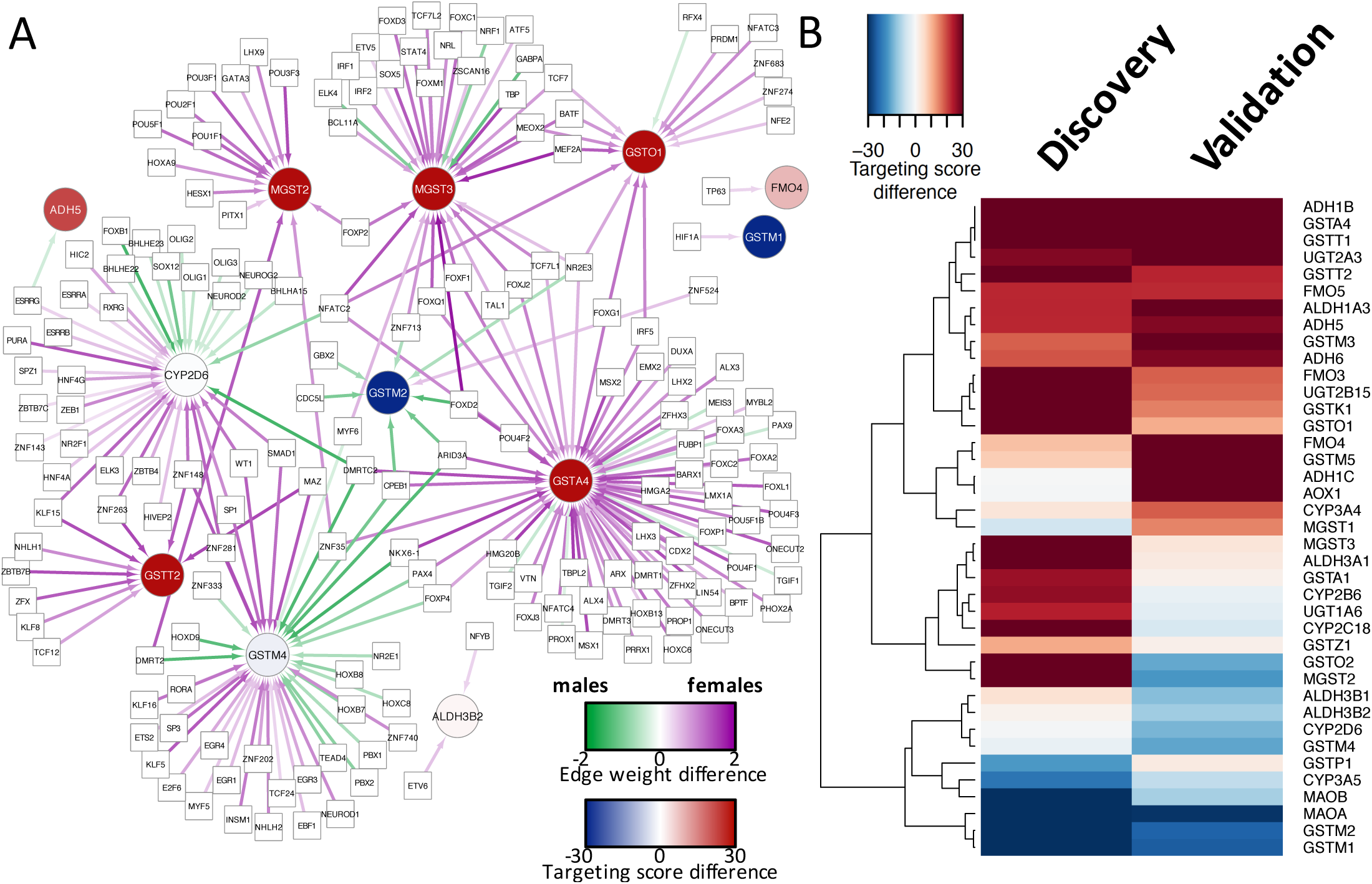
Differential targeting of genes in the drug metabolism pathway. A) Subnetwork representation of the edges in the drug metabolism pathway that are most significantly different between male and female regulatory networks (FDR<0.05). B) Heatmap of gene targeting score difference for all the analyzed genes in the drug metabolism-cytochrome P450 pathway for the discovery and validation dataset analyses. For visualization purposes, the color scale was saturated at 30. Positive values indicate higher levels in females, and negative values indicate higher levels in males.

Next, we confirmed the drug metabolism pathway differential targeting in an independent dataset. As previously described, our validation dataset was obtained from GEO and included 1,193 samples from five independent studies. We repeated the pipeline described above to reconstruct single-sample gene regulatory networks and to perform differential targeting analysis in the validation dataset. The pathways more strongly targeted in males did not reach statistical significance in this validation dataset (Table S2). However, we again identified the drug metabolism-cytochrome P450 (FDR=0.009), and the metabolism of xenobiotics by cytochrome P450 (FDR=0.012) pathways as enriched for genes that were more strongly targeted in females.

Some known tumor molecular features, such as CpG island methylator phenotype (CIMP), chromosomal instability (CIN) phenotype, and mismatch repair (MMR) deficiency, may be sex-biased and/or associated with disease prognosis and treatment response. Since part of the validation group was profiled for these features, we repeated the differential targeting analysis for the validation group adjusting for CIMP, CIN, and MMR. Again, we confirmed females have higher targeting of the drug metabolism-cytochrome P450 (FDR=0.032), and the metabolism of xenobiotics by cytochrome P450 (FDR=0.038).

This validation in an independent dataset, and one derived using a different technology (microarrays rather than RNA-Seq), gives us a high degree of confidence in the observed sex-specific regulation of drug metabolism. Indeed, the genes in the drug metabolism pathway with higher gene targeting score difference are highly consistent between the discovery and validation datasets (Figure 3B). The main regulatory sex-differences are associated with genes important during catabolism and detoxification, for example *ADH1B, GSTA4*, and *FMO5*.

We also analyzed differential targeting of the drug metabolism pathway between males and females in healthy colon tissue. We compared healthy colon tissue samples from 223 males and 153 females (obtained from Chen et. al. (22)) by performing differential targeting analysis after adjusting for age and race. We found no significant difference for the drug metabolism-cytochrome P450 pathway (FDR=0.31), nor the metabolism of xenobiotics by cytochrome P450 (FDR=0.44) in healthy tissues. This indicates the sex-differences observed in the regulatory patterns of the drug metabolism pathway only become apparent in colon cancer but not in healthy tissue.

### Higher targeting of the drug metabolism pathway is associated with better overall survival in females treated with adjuvant chemotherapy

Considering that treatment benefit and survival outcome are largely dependent on tumor stage, we performed a survival analysis to evaluate chemotherapy treatment benefit in each stage. Only patients with stage and treatment information were selected for these survival analyses (n=514 from GSE39582). Stage I patients were excluded because none of these patients were treated with chemotherapy. For stage II, there was no overall survival difference between patients with and without chemotherapy treatment (Figure S2). This was expected, as many studies show only a small or no benefit from chemotherapy for stage II patients, and only a small subgroup of patients with more aggressive tumors benefits from treatment (40). Chemotherapy treatment benefit was clear for stage III patients: treated patients had a significantly higher overall survival probability than non-treated patients. For stage IV, we did not find a survival difference after chemotherapy treatment, possibly due to the small number of samples included in this analysis (n=45). Therefore, we limited subsequent survival analysis to stage III patients (n=190).

To evaluate whether the strength of targeting for drug metabolism pathway genes is associated with outcome, we performed overall survival analysis after stratifying patients by how strongly the pathway was targeted. For this, patients in the validation dataset were divided in two groups of equal size (named low and high targeting groups) based on the median targeting score of genes in the drug metabolism pathway. For patients not treated with adjuvant chemotherapy, targeting of the drug metabolism pathway did not change the overall survival probability, showing that high targeting is not a prognostic factor (Figure 4A).

**Figure 4.**
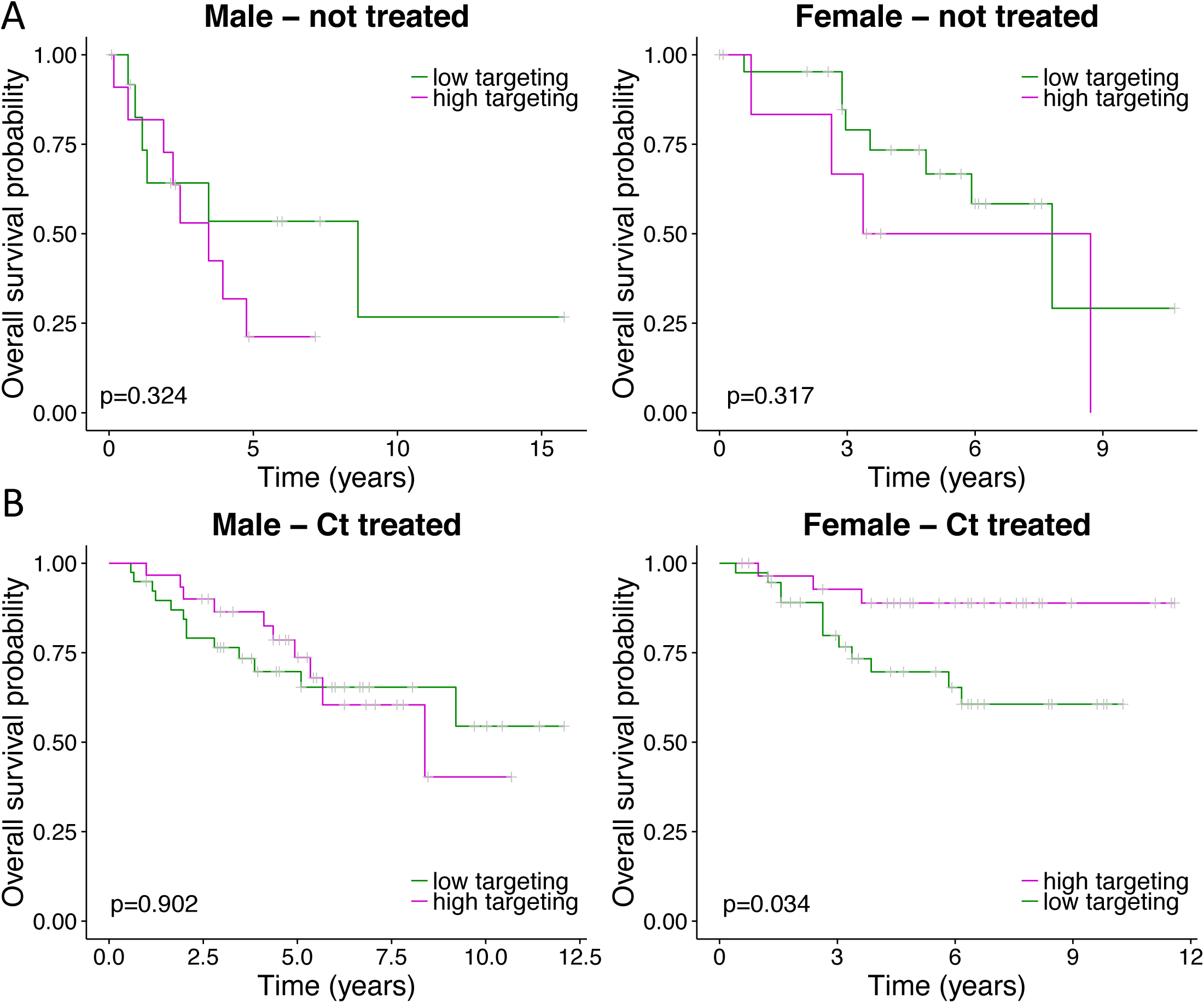
Overall survival analysis based on targeting of the drug metabolism pathway. Kaplan-Meier curve of stage III colon cancer patients not treated (A) or treated (B) with adjuvant chemotherapy. Patients were stratified by low and high targeting groups based on the median gene targeting score across all genes in the drug metabolism pathway. P-values were computed using the log-rank test to evaluate the overall survival risk between the low targeting (green) and high targeting (purple) subgroups of patients.

Next, we assessed the clinical impact of the drug metabolism pathway targeting for patients treated with adjuvant chemotherapy. Consistent with the weaker targeting of the drug metabolism pathway observed in male networks, we found that gene targeting strength of the drug metabolism pathway did not change the overall survival probability for males receiving adjuvant chemotherapy (Figure 4B). In contrast, in females who received treatment, higher targeting of the drug metabolism pathway was associated with a better overall survival. The 10-year overall survival probability was 61% (95% CI: 45%-82%) for females with low targeting and 89% (95% CI: 78%-100%) for females with high targeting (log-rank test p=0.034). There was no statistically significant difference between the females high and low targeting groups for the available covariates (age, MMR, CIMP, and CIN).

## Discussion

The use of standard differential expression analysis between males and females have found the expected differential regulation of sex-chromosome genes (41–43). However, differential targeting analysis identified sex-specific differences are also driven by global transcriptional regulatory processes, a phenomenon we had seen previously in chronic obstructive pulmonary disease (COPD) (15).Using a network-based approach we found that not only are genes associated with drug metabolism differentially regulated in males and females, but also that patterns of gene targeting are associated with clinical outcome, particularly in women. Specifically, we find that greater targeting of the drug metabolism pathway is associated with a better overall survival in females treated with adjuvant chemotherapy.

Many of the genes associated with the drug metabolism pathway are highly polymorphic, and they have been associated with colon cancer risk (44–46). For example, a meta-analysis showed that *GSTM1* and *GSTT1* null genotypes increase the risk of colorectal cancer in Caucasian populations (47). For lymphoblastoid cell lines (LCLs), it has been shown that the KEGG pathway “metabolism of xenobiotics by cytochrome P450” is enriched for genes more expressed in females compared to males (48). In healthy colon tissues, we found no significant sex-differences in the regulatory patterns of the drug metabolism pathway. The differences only became evident in colon cancer tissues, and these differences are independent of the disease stage.

The genes with the largest sex differences belong to the glutathione S-transferase family and they are highly targeted in females. GSTs are metabolizing enzymes that play a key role during neutralization and elimination of toxic compounds and xenobiotics (38). Thus, considering the role of the differentially targeted genes found on samples before treatment, one can expect that each sex might respond differently to chemotherapeutic treatment, and ultimately have a different survival outcome.

We note the analyzed data might be influenced by cellular heterogeneity or clinical heterogeneity and risk factor profiles (such as dietary habits, and family history). To reduce confounding effects, the data was corrected for known covariates, such as age, race, and disease stage. Although our study has a small number of samples treated with adjuvant chemotherapy and with survival information, the identification of differential regulation of drug metabolism pathways in independent datasets provides support for our conclusions. This suggests that clinical trials and other experiments should carefully consider the manifestation of sex differences and be statistically powered to understand the impact of sex in outcomes.

Even though there are significant sex differences regarding risk, prognosis, treatment response, and chemotherapeutic toxicity related to colon cancer, management of colon cancer is not based on sex, and the molecular features that drive the sex differences are still poorly understood. We found that colon cancer has significant sex differences related to TFs regulatory patterns. Most importantly, targeting of the drug metabolism pathway was predictive of higher overall survival in women who received adjuvant chemotherapy. The determination that sex-specific targeting could discriminate between long term and short term survivors raises the possibility of using gene regulatory network analyses in other diseases. Indeed, the regulatory network analysis method described here can easily be used to understand how sex influences progression and response to therapies in other cancer types and complex diseases and may help motivate development of sex-specific approaches to disease treatment.

